# Molecular glue CELMoD compounds are allosteric regulators of cereblon conformation

**DOI:** 10.1101/2022.07.02.498551

**Authors:** Edmond R. Watson, Scott J. Novick, Mary E. Matyskiela, Philip P. Chamberlain, Andres Hernandez de la Peña, Jin-Yi Zhu, Eileen Tran, Patrick R. Griffin, Ingrid E. Wertz, Gabriel C. Lander

## Abstract

Cereblon (CRBN) is an ubiquitin ligase (E3) adaptor protein co-opted by CRBN E3 Ligase Modulatory Drugs (CELMoD) agents that target therapeutically-relevant proteins for degradation. Prior crystallographic studies defined the drug-binding site within CRBN’s Thalidomide Binding Domain (TBD), but the allostery of drug-induced neosubstrate binding remains unclear. We therefore performed cryo-EM analyses of the DDB1∼CRBN apo-complex, and compared these structures with DDB1∼CRBN in the presence of CELMoD compounds alone and complexed with neosubstrates. Association of CELMoD compounds to the TBD is necessary and sufficient for triggering CRBN allosteric rearrangement from an “open” conformation to the canonical “closed” conformation. Importantly, the neosubstrate Ikaros only stably associates with the closed CRBN conformation, illustrating the importance of allostery for CELMoD compound efficacy, and informing structure-guided design strategies to improve therapeutic efficacy.

## Introduction

Eukaryotic proteins are targeted for degradation through covalent attachment of ubiquitin moieties to target protein residues, primarily lysine side-chains^1^. Ubiquitination is facilitated by ubiquitin ligases (E3s), which coordinate and position targeted substrates for ubiquitin ligation, often through adaptor proteins and interchangeable receptor modules. Substrate specificity of one such E3 complex, the Cullin-4 RING ligase (CRL4), is mediated by the cereblon (CRBN) substrate receptor. CRBN facilitates the transfer of ubiquitin to substrate proteins by reversibly interacting with the CRL4 core complex, which consists of the Cullin-4 scaffold, the DDB1 adaptor protein, and RING finger protein ROC1 (**Supplementary Fig. 1A**). CRBN is also the cellular receptor for a class of drugs known as CELMoD agents, which bind to a conserved hydrophobic pocket in CRBN to create a molecular surface capable of recruiting “neosubstrates” – cellular proteins that are targeted only in response to drug. Such ligand-induced CRL4-CRBN recruitment facilitates neosubstrate ubiquitination and subsequent proteasomal degradation, thereby dictating therapeutic efficacy^2^. The CELMoD agent lenalidomide (Revlimid) has been used as a first-line therapy for multiple myeloma and other hematological malignancies for more than a decade, and next-generation CELMoD compounds markedly improve patient outcomes in clinical trials. Although crystallographic structural snapshots of CRBN∼DDB1 bound to a variety of drugs and neosubstrates have facilitated a molecular description of the stable quaternary complex, the allostery associated with substrate targeting remains unknown and the structural features of the unliganded complex has yet to be described.

CRBN is a 50 kiloDalton protein containing three folded domains: an N-terminal Lon protease-like domain (hereafter Lon domain), an intermedial helical bundle (HB), and a C-terminal Thalidomide-Binding Domain (TBD) that harbors the drug binding pocket. CRBN is recruited to the Cullin-4/ROC1 ligase module *via* the adaptor protein DDB1, which consists of three WD-40 beta propeller domains (BPA, BPB, BPC). The HB of CRBN docks into a central hydrophobic cleft formed at the interface of BPA and BPC of DDB1, while the mobile BPB domain interacts with Cullin-4, positioning CRBN-bound substrates for ubiquitination. Over a dozen crystal structures of the CRBN∼DDB1 complex show the Lon domain and TBD of CRBN tightly interacting with one another, that can be described as a “closed” conformation (CRBN^closed^)^2-9^. These structures of the closed conformer bound to small molecules have provided the basis for structure-guided drug design of next-generation CELMoD agents with enhanced neosubstrate specificity and potency, such as CC-92480 (mezigdomide).

While it is thought that association of the TBD and Lon domains are concomitant with drug and neosubstrate binding, this assumption is challenged by structures of the isolated TBD bound to ligands, demonstrating that the TBD is competent for drug binding in the absence of the Lon domain^10^. Further, two crystal structures have captured the Lon domain and TBD in an “open” conformation (CRBN^open^) where the two domains are separated and positioned at a ∼45° angle relative to one another. These reports raise questions about the assumed constraints associated with drug accessibility and binding, as well as the mechanisms by which ligands modify the surface chemistry of CRBN to enable neosubstrate recruitment and subsequent ubiquitination and degradation^3,4^. If CRBN^open^ represents a physiological conformer (i.e. not an artifact of crystallization) we must reconsider recruitment of drug and neosubstrates in the context of these two conformers. Additionally, the dynamics of interconversion between the open and closed conformers, the effect that ligands and neosubstrates have on this process, and how these dynamics affect neosubstrate positioning and subsequent degradation may have profound effects on the design, and ultimately the therapeutic impact, of next-generation CELMoD compounds.

### CRBN basally adopts an open conformation when bound to DDB1

To better understand the spectrum of CRBN conformations, we used cryo-electron microscopy (cryo-EM) to examine the population ensemble of CRBN∼DDB1 conformers present in solution under various conditions. To overcome preferred orientation and partial denaturation/dissociation of the complex during cryo-EM grid preparation, a combination of grid pretreatment with a CRBN-agnostic protein, incorporation of mild amphiphilic detergent, and sample dilution immediately prior to plunging improved particle homogeneity and distribution in ice (see Methods). Unexpectedly, whether copurified with either full-length DDB1 or DDB1 lacking BPB (CRBN-DDB1^ΔBPB^, a construct commonly used for X-ray crystallography), apo-CRBN exclusively adopts an open conformation, with the TBD and Lon domains separated from one another (**Fig. 1A, Supplementary Fig. 1B**).

**Figure 1.**
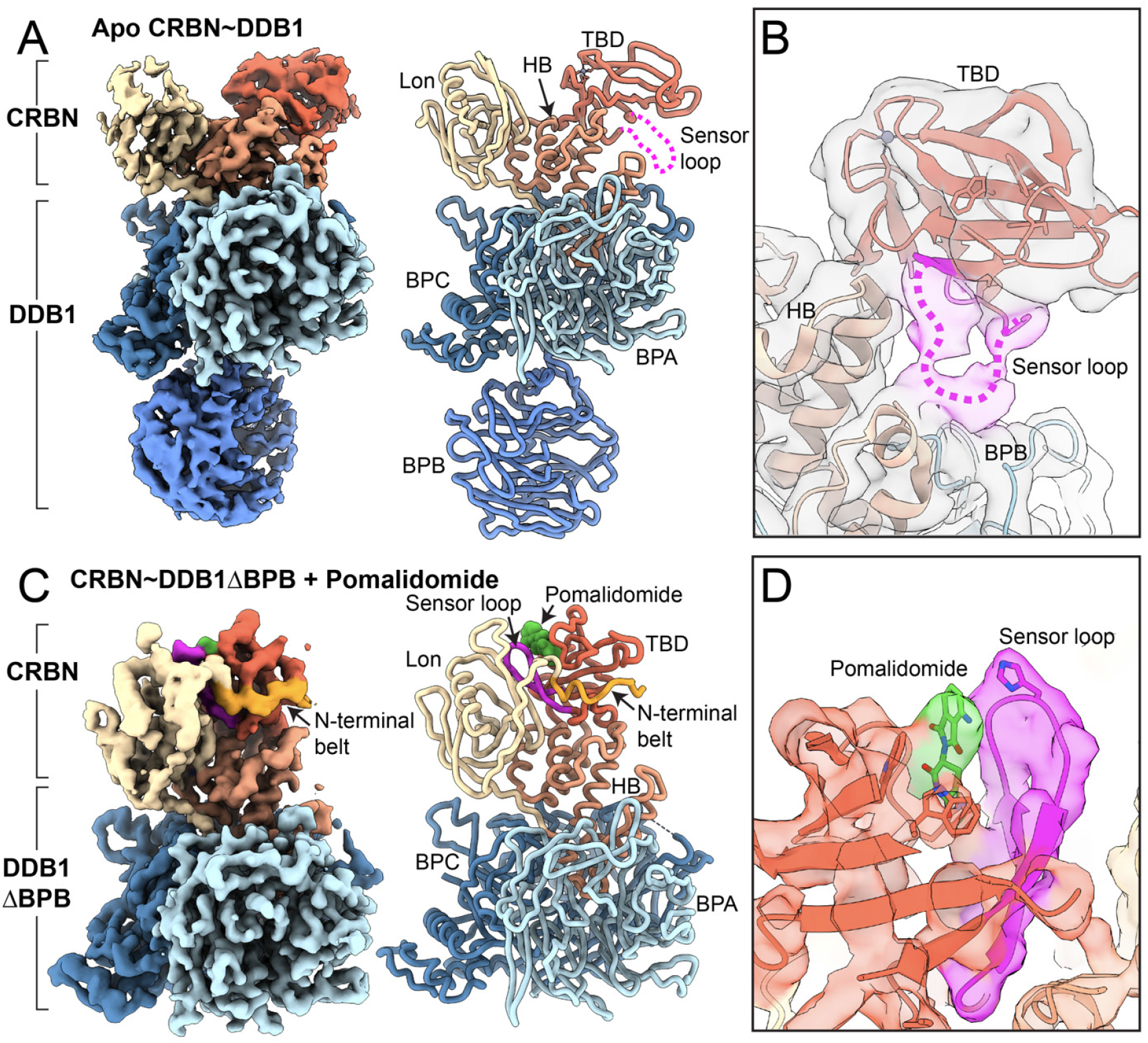
CRBN^open^ is allosterically modulated to CRBN^closed^ by pomalidomide. **(A)** ∼3.5 Å resolution cryo-EM reconstruction of CRBN∼DDB1 in the unliganded apo form. The Lon domain (tan) is separated from the TBD (red), while the helical bundle (HB, light red) mediates interaction with DDB1, composed of beta propeller A (BPA, light blue), beta propeller B (BPB, medium blue), and beta propeller C (BPC, steel blue). CRBN and DDB1 are thresholded independently to illustrate important features. To the right, a ribbon representation of the CRBN DDB1 complex modelled from density is shown, colored as in (A). The sensor loop is denoted as a dotted magenta line extending away from the TBD domain. **(B)** The unsharpened cryo-EM map shows the path of the sensor loop, which interacts with the HB and BPC of DDB1. **(C)** Surface representation of the ∼3.9 Å resolution cryo-EM reconstruction of CRBN DDB1 in the closed form in complex with pomalidomide. The N-terminal belt (orange) is observe wrapping around the TBD domain. To the right, a ribbon representation of CRBN∼DDB1^ΔBPB^ in complex with pomalidomide is shown, with the sensor loop (magenta) adopting a beta-hairpin organization that is tightly packed between the TBD and Lon domains. **(D)** A detailed view of the density corresponding to pomalidomide (green) and sensor loop (magenta) in CRBN^closed^.

Our ∼3.5 Å resolution cryo-EM structure confirms the physiological existence of the open conformer, and provides the first opportunity to examine this CRBN conformation in the absence of ligand or substrate. Notably, we do not observe density for the “sensor loop,” a beta-insert hairpin within CRBN’s TBD (residues ∼341-361) that has been shown to directly bind IMiDs and neosubstrates in prior structures^2,11^, at its canonical position adjacent to the thalidomide binding pocket. Rather, we observe strong, low-resolution, density corresponding to the sensor loop engaged in previously unobserved interactions with a helix in CRBN’s HB (residues ∼210-220) and a loop of BPB in DDB1 (residues ∼776-780) (**Fig. 1B**). These regions of the HB and DDB1 are usually disordered in crystal structures^4,9,11,12^, consistent with the flexibility of this region observed in our cryo-EM density. However, the clearly-observed association between the sensor loop and this HB/DDB1 interface in our cryo-EM structures suggests that interaction between these flexible loops plays a role in maintaining the TBD in the CRBN^open^ conformer.

### Binding of CELMoD compounds drives conformational rearrangement within CRBN

To better understand how CELMoD agents interact with and influence this open CRBN conformation, we added saturating amounts of different compounds to CRBN-DDB1 for cryo-EM analyses. Addition of pomalidomide was sufficient to induce conformational rearrangement within CRBN for only ∼20% of the particles. The ∼3.9 Å resolution structure of CRBN^closed^ arising from pomalidomide-bound particles shows the ∼15kDa TBD positioned adjacent to Lon domain, and unambiguous density within the ligand-binding pocket consistent with pomalidomide association (**Fig. 1C**). Necessarily, the sensor loop in CRBN^closed^ is observed in the canonical beta-hairpin fold interfacing the TBD and Lon domain, and a portion of the CRBN N-terminus (residues 48-63) that is disordered in the CRBN^open^ becomes ordered, extending from the Lon domain, trussing the TBD and supporting the closed conformation (**Fig. 1D**). To better define the role of these N-terminal residues, hereafter referred to as the N-terminal belt, in CRBN closure, we truncated 63 residues from the CRBN N-terminus (CRBN^ΔNTD^) and incubated the CRBN^ΔNTD^∼DDB1 with saturating pomalidomide concentrations. Notably, deletion of the N-terminus all but abolishes CRBN closure, with less than ∼2% of the complexes adopting a closed conformer with a poorly ordered sensor loop, underscoring the importance of the CRBN N-terminal belt in stabilizing the CRBN^closed^ conformer (**Supplementary Fig. 2A**).

While we expected to observe the pomalidomide ligand in the binding pocket of only the CRBN^closed^ conformer, we were surprised to find strong density corresponding to pomalidomide within the binding pocket of CRBN^open^ as well (**Supplementary Fig. 3A**). Furthermore, even with ligand bound, and in stark contrast to prior crystal structures of the CRBN^open^ conformer, the sensor loop of our cryo-EM CRBN^open^ conformer remains engaged in interactions with the CRBN HB and DDB1 as we observe in the apo-CRBN conformer (**Supplementary Fig. 3B**). These results indicate that limitations in neosubstrate recruitment, that directly translate to drug efficacy, may not solely correlate with CRBN-binding kinetics. Rather, conversion of the CRBN^open^ to the CRBN^closed^ conformation may be a key mechanistic bottleneck in the allosteric transformation of apo-CRBN to a neosubstrate-binding conformation following ligand association.

We were intrigued by the observation that saturating concentrations of pomalidomide did not induce widespread closure of the CRBN domains, despite high occupancy. We rationalized that next-generation CELMoD agents with improved binding affinity may also have a more pronounced allosteric effect through enhanced interactions with the sensor loop, contributing to adoption of the sensor loop to the CRBN^closed^–competent β-hairpin conformation. Indeed, analyzing CRBN∼DDB1 in the presence of CC-220 (iberdomide), a molecule with ∼20-fold improved affinity and ∼24-fold enhanced Ikaros degradation^12^, shows nearly 50% of particles have shifted to the CRBN^closed^ conformer (**Fig. 2A**). As with pomalidomide, with excess ligand present, we observed density consistent with iberdomide within the binding pocket of both the CRBN^closed^ and CRBN^open^ conformers, suggesting drug occupancy is not responsible for the increased allosteric transition. Notably, a prior crystal structure of the iberdomide-bound CRBN^closed^ shows that the additional phenyl and morpholino moieties of the iberdomide molecule arch over the top of the sensor loop β-hairpin^12^, likely aiding in the stabilization of this structural motif. These data support our proposed allosteric model whereby enhanced ligand interaction with the sensor loop stabilizes the β-hairpin conformation, which promotes the closure of the TBD and Lon domains and ultimately translates to improved neosubstrate ubiquitination and subsequent degradation efficacy.

**Figure 2.**
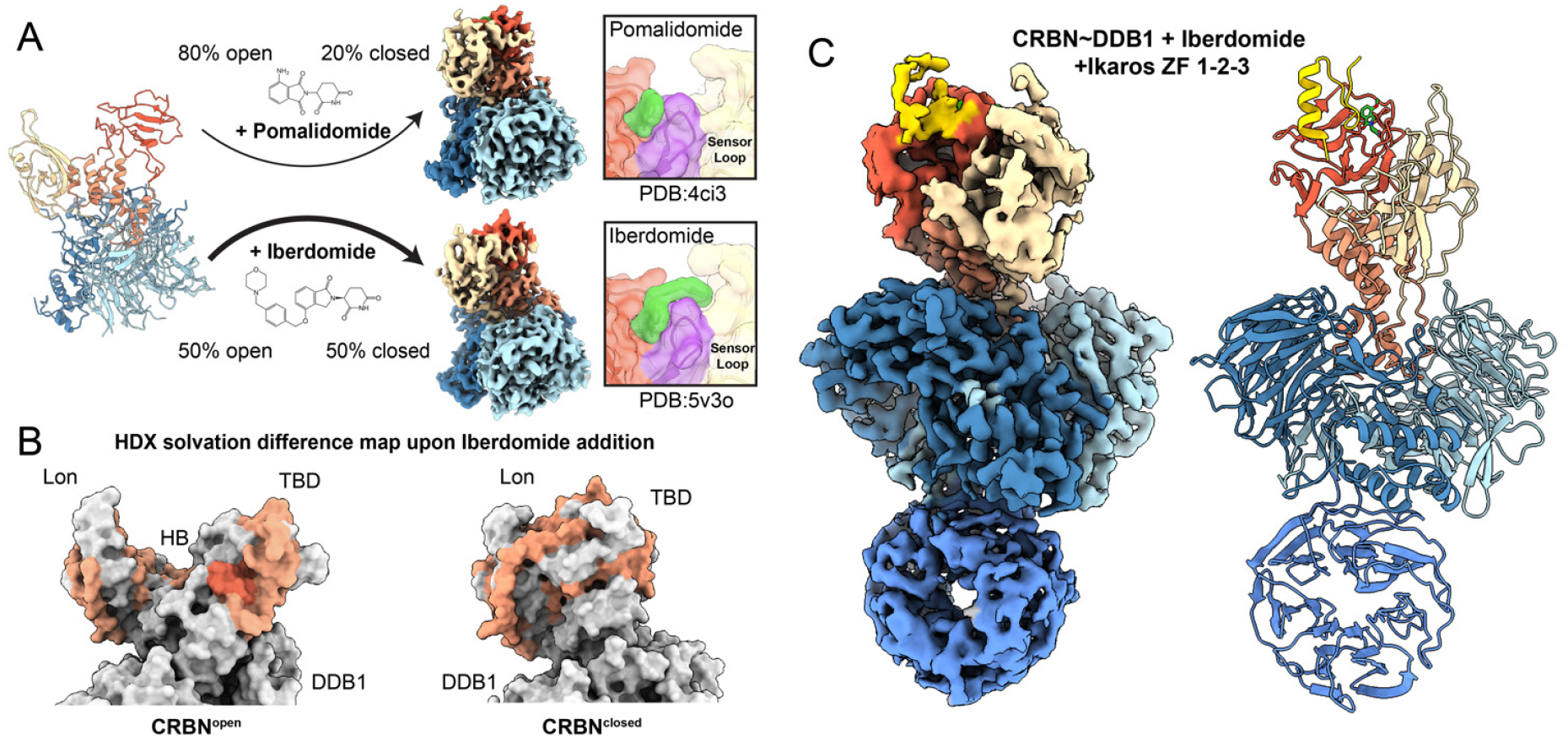
Improved CELMoD compounds drive CRBN^closed^ and recruit Ikaros. **(A)** The CRBN^open^ transition to CRBN^closed^ is differentially regulated between pomalidomide (∼20% particles adopt CRBN^closed^) and the CELMoD agent iberdomide (∼50% particles adopt CRBN^closed^). Upper right, the ∼3.9 Å resolution cryo-EM structure of pomalidomide-induced CRBN^closed^. Lower right, the ∼3.7 Å resolution reconstruction of iberdomide-induced CRBN^closed^. The inset panels on the far right depict surface representations of published crystal structures illustrating the ligand (green) interactions with sensor loop (magenta). **(B)** Colored space-filling representation of CRBN models with residue-specific coloring according to changes in solvency upon addition of iberdomide as detected by HDX-MS. Orange residues correspond to peptides with mild HDX differential, red residues correspond to peptides with large HDX differential. Mixed conformations are expected during the experiment, so CRBN^open^ and CRBN^closed^ are both shown. **(C)** The ∼3.6 Å resolution reconstruction of iberdomide-induced CRBN^closed^ in complex with Ikaros ZF1-2-3 (gold). DDB1 and CRBN are thresholded independently to illustrate important features.

In order to further investigate the allosteric influence of Iberdomide on CRBN closure, we used hydrogen-deuterium exchange mass spectrometry (HDX-MS) to profile changes in solvent accessibility of CRBN∼DDB1 peptides in the presence of iberdomide. Compared with unliganded CRBN∼DDB1, addition of drug substantially reduced solvation for regions within the Lon, TBD, and HB domains found at the domain-domain interface of the closed conformer, consistent with transition from CRBN^open^ to CRBN^closed^ (**Fig. 2B, Supplementary Fig. 4A**), while the DDB1 surfaces were unchanged (**Supplementary Fig. 4B**). As expected, the most extreme change is observed for the sensor loop itself, consistent with formation of the buried hairpin we observe in CRBN^closed^ (**Fig. 1C**). These findings offer a potential new modality for drug development, wherein properties influencing not only binding kinetics for CRBN and neosubstrates^13^, but also the capacity to stabilize the β-hairpin conformer of the sensor loop as a means of both initiating and maintaining the closed conformation should be considered. The CRBN^closed^ conformation may in turn be a prerequisite for substrate recruitment.

Indeed, the substantial population of ligand-bound CRBN^open^ conformers in our datasets indicates that ligand binding is not concurrent with adoption of the CRBN^closed^ conformer, and that CRBN closure is dependent on subsequent allosteric triggers. We posit that ligand binding is involved in inducing the sensor loop’s adoption of the beta-hairpin arrangement, thereby untethering the TBD from the CRBN HB to enable the CRBN^closed^ conformer for substrate interaction. Despite extensive focused classification of the tether region, we were unable to identify a population of liganded CRBN^open^ conformer where the sensor loop is untethered from the HB in any of our datasets. This finding indicates that ligand-mediated detachment of the sensor loop from the HB is concomitant with adoption of the structured beta-hairpin and triggers immediate closure of the CRBN TBD and Lon domains.

### CELMoD compounds mediate recruitment of Ikaros to the closed conformation

We next aimed to probe the role of drug binding in neo-substrate recruitment and positioning in the context of the CRBN^open^ and CRBN^closed^ conformers. Prior crystallographic studies yielded structures of CRBN^open^ conformers bound to neosubstrates^3,8^, although it was speculated that these conformations may have resulted from crystallization conditions and lattice contacts. To investigate drug-mediated CRBN∼DDB1 recruitment of neosubstrates, we employed the zinc-finger transcription factor Ikaros, a cellular target of pomalidomide and iberdomide. We generated different constructs containing tandem Ikaros zinc finger (ZF) 1-3 domains (**Supplementary Fig. 3C**) for increased recruitment efficiency^3^ and incubated each individually with CRBN∼DDB1 in the presence of pomalidomide or iberdomide. Regardless of drug identity or Ikaros ZF construct design, our single-particle analyses reveal density consistent with the location and positioning of a single Ikaros zinc finger motif associated with the TBD domain of the CRBN^closed^ conformer (**Fig. 2C, Supplementary Fig. 3D**), again confirmed in HDX-MS analyses showing even more pronounced changes for residues within the sensor loop that contact the known Ikaros recruitment motif (**Supplementary Fig. 4C-D**). Notably, we were unable to detect any Ikaros association with the CRBN^open^ conformer, consistent with a mechanism whereby drug-induced CRBN closing occurs prior to substrate recruitment. This further demonstrates the importance of an allosteric control mechanism, whereby drug-induced CRBN closing occurs prior to neosubstrate recruitment.

### Beta propeller B of DDB1 alternates between three distinct conformations

The Cullin-4∼ROC1 ligase core recruits the DDB1∼CRBN^closed^-Ikaros complex by associating with the second beta-propeller of DDB1 (called BPB), which is located distally to CRBN. BPB extends from the DDB1 core as a mobile element, and this flexibility has been implicated in the mechanism of DDB1 as a promiscuous adaptor that bridges Cullin-4 with structurally diverse substrate-binding modules. Correspondingly, BPB has been captured in a variety of conformations in several prior studies^14^. We sought to describe the motion of BPB and its potential relevance to Ikaros positioning for ubiquitination in the context of the CRL4-CRBN complex assembly. Intriguingly, our image analyses of the BPB domain within our apo CRBN data reveals that, rather than demonstrating a full continuum of motion, the BPB position is limited to three metastable positions. Approximately 65% of the particles adopts a “linear” orientation, in which the broad face of BPB is in a similar orientation as BPC (**Fig. 3A, left**). A ∼70° rotated “hinged” orientation, where the BPB face more closely matches BPA is the second major class observed, with ∼20-30% of particles (**Fig. 3A, middle**), and only a minor population contain BPB in the third, “twisted” orientation, rotated ∼140° from the linear orientation (**Fig. 3A, right**). Further classification of particles contributing to each of these states reveals only minor positional variability for BPB, as has been previously described^14^, centered around these discrete conformations.

**Figure 3.**
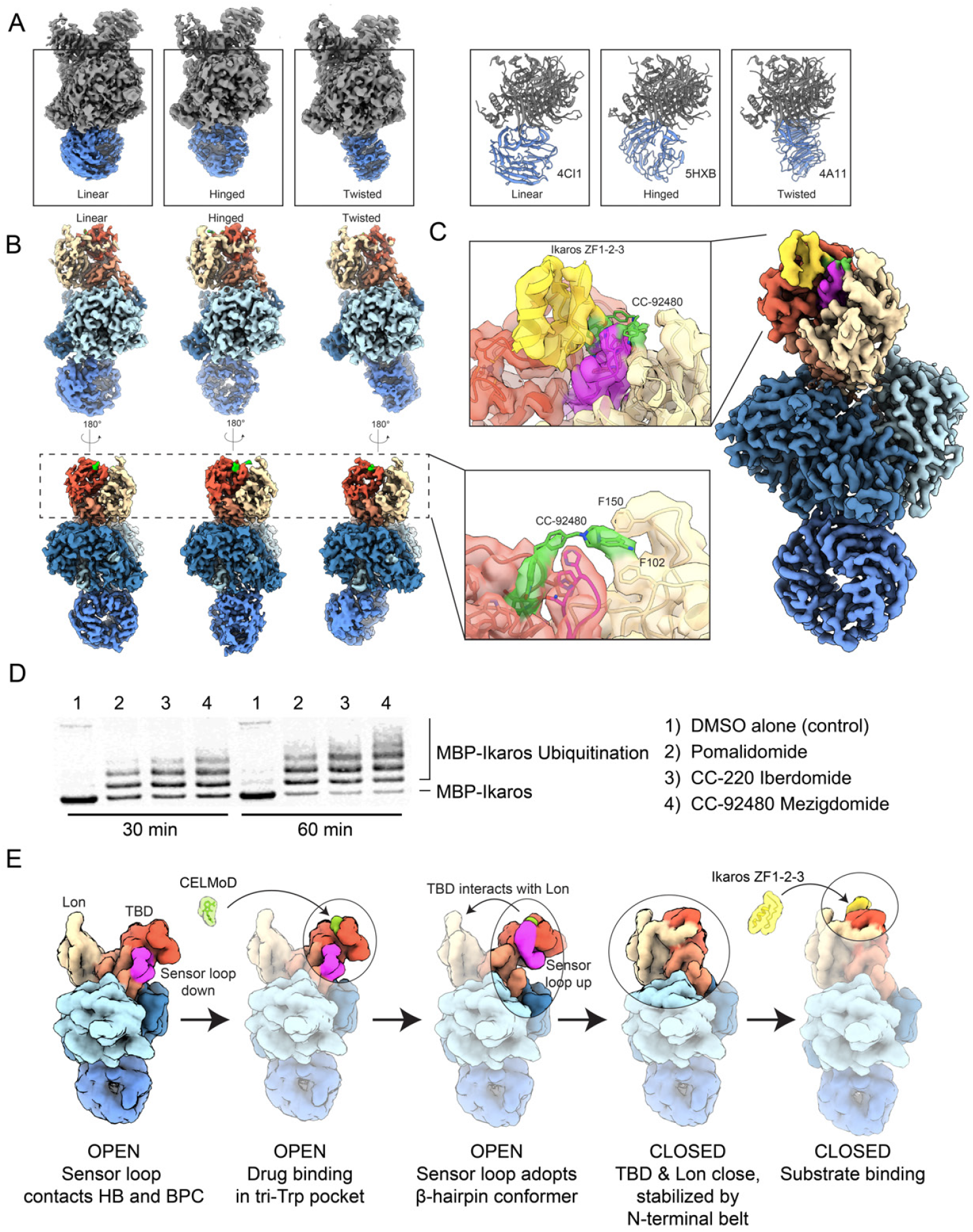
DDB1 and next-generation CELMoD agents further poise CRBN substrates for ubiquitination in disease contexts. **(A)** 3D classification of unliganded CRBN∼DDB1 yields three discrete positions of DDB1’s mobile BPB propeller (colored blue). Left, ∼3.5 Å resolution cryo-EM reconstructions of CRBN∼DDB1 with BPB in linear (left), hinged (middle) or twisted (right) position. Right, structural models in similar orientations for reference, adopting linear (PDB: 4CI1, left), hinged (PDB: 5HXB, middle), and twisted (PDB: 2HYE, right). **(B)** The ∼3.2 Å resolution cryo-EM reconstructions of CRBN∼DDB1 complexed with mezigdomide with linear (left), hinged (middle), and twisted (right) positions of BPB. Inlay, ∼3.1 Å resolution focused refinement of particles from all three orientations reveal a connection between mezigdomide and Lon domain, “stapling” the CRBN^closed^ conformer. **(C)** The ∼3.1Å resolution reconstruction of mezigdomide-induced CRBN^closed^ bound to Ikaros ZF1-2-3. The inlay shows a focused view of connection between mezigdomide and Lon domain, stapling CRBN^closed^ in the presence of Ikaros. **(D)** *in vitro* ubiquitination assay following MBP-Ikaros ubiquitination in the presence of varying molecules used in this study, labeled on the right. **(E)** Mechanistic model illustrating pathway of assembly. First, unliganded CRBN is open with the sensor loop attached to HB. Second, ligand is added and associates with TBD in the hydrophobic pocket. Third, binding of ligand in the tri-Trp pocket stabilizes the sensor loop as a beta-hairpin detached from the HB. Fourth, with sensor loop refolding promotes transition of CRBN^open^ to CRBN^closed^ without an observed intermediate, and N-terminal belt is seated (orange). Fifth, substrate is recruited to the CRBN^closed^ for subsequent ubiquitination by CRL4.

These three principal orientations were observed in recent low-resolution cryo-EM structures of DDB1-DCAF1 bound to Cullin-4^15^, and our structures demonstrate that these BPB positions are intrinsic to DDB1, irrespective of the substrate adaptor module or interaction with Cullin. Although we were unable to discern allosteric coordination upon ligand- or Ikaros-binding, the population distribution of the states vary significantly compared to those observed for the Cullin-bound DDB1^15^. Notably, the “twisted” orientation that we observe in less than 10% of our particles is the majority conformer when bound to Cullin^15^, suggesting that association with Cullin may substantially modulate BPB positioning between these three states, thereby influencing proximity and positioning of neosubstrates for ubiquitination.

### Implications for Relapsed/Refractory Multiple Myeloma

Given the potential clinical relevance of the observed CRBN allostery, we next used our method to characterize the mechanisms of action responsible for the improved efficacy of next-generation CELMoD molecules. Mezigdomide, for example, was recently shown to display efficient, rapid neosubstrate degradation kinetics and is considerably more efficacious in the treatment of relapse/refractory patients who no longer respond to primary treatment with lenalidomide and/or pomalidomide^16^. To investigate the structural features responsible for the enhanced efficacy of this compound, we incubated mezigdomide with CRBN∼DDB1 for cryo-EM analyses, and were astonished to find that 100% of the complexes adopted the CRBN^closed^ conformer. Density corresponding to mezigdomide is clearly observed in the canonical binding site in the ∼3.2 Å structures of CRBN-DDB1 with BPB in the three discrete orientations (**Fig. 3B**). Importantly, we observe protruding drug density that extends to the distal Lon domain, a feature that has not yet been observed in the structures of any other liganded CRBN (**Fig. 3B, right**). We were able to unambiguously fit mezigdomide into the density to reveal that the benzonitrile moieity is likely stabilized by aromatic or cation-pi interactions with CRBN Phe-102 and Phe150. Association of both the glutarimide and the benzonitrile moieities to distinct domains within CRBN both provides avidity for improved drug binding and serves as a molecular staple of the closed conformation, which is traditionally accomplished by the CRBN N-terminal belt (**Fig. 1C**). Ikaros is recruited to mezigdomide-bound CRBN∼DDB1 (**Fig. 3C**) and its ubiquitination efficiency is enhanced in assays with Cul4∼Roc1 (**Fig. 3D**). Intriguingly, within the mezigdomide-bound dataset we observe a substantial population of CRBN^closed^ particles that lack discernible density for the stabilizing N-terminal belt (**Supplementary Fig. 4B**). This indicates functional redundancy in securing the closed conformer, and suggests that mezigdomide further promotes neosubstrate recruitment and subsequent ubiquitination by prolonging the mechanistic cycle of closure, neosubstrate binding, N-terminal engagement, and stepwise reversal to the open conformation. These data elucidate an important new compound design modality and describe the structural mechanism for enhanced therapeutic benefit of mezigdomide in patients unresponsive to pomalidomide, further illustrating the therapeutic value of allosteric control.

Taken together, our results provide novel mechanistic insights that elucidate the therapeutic efficacy of CELMoD compounds, a class of molecules that epitomize molecular glues and are central players in the field of targeted protein degradation. In so doing, we highlight previously unappreciated allosteric aspects for consideration in the design of CRBN-directed molecular glue therapeutics (**Fig. 3E**). Through characterization of the conformational rearrangements inherent to the CRBN∼DDB1 system, and how three distinct degrader molecules impact allostery and neosubstrate-binding capacity, we have shown how conformational control of the mobile drug-binding TBD within CRBN has cryptically driven the therapeutic success of neosubstrate targeting agents. These findings provide an important context for interpreting physiological roles of endogenous CRBN, and serve as the basis for rational development of therapeutic compounds that promote and stabilize the CRBN^closed^ conformer to enhance therapeutic efficacy. Furthermore, by highlighting the critical rearrangements and interactions that are required for CRBN closure, we yield insights into the allosteric perturbations that may be introduced by patient mutations throughout the *CRBN* gene, and more importantly how they may be overcome via rational drug design.

## Materials and Methods

### Protein expression and purification

ZZ-domain-6xHis-thrombin-tagged human CRBN (amino acids 5-442) and full-length human DDB1, or DDB1^ΔBPB^ which removes residues 396-706 and adds a GNGNSG linker, were co-expressed in SF9 insect cells in ESF921 medium (Expression Systems), in the presence of 50 µM zinc acetate. Cells were resuspended in buffer containing 50 mM Tris-HCl (pH 7.5), 500 mM NaCl, 10 mM imidazole, 10% glycerol, 2 mM TCEP, 1X Protease Inhibitor Cocktail (San Diego Bioscience), and 40,000 U Benzonase (Novagen), and sonicated for 30 s. Lysate was clarified by high speed centrifugation at 108,800 *g* for 30 min, and clarified lysate was incubated with Ni-NTA affinity resin (Qiagen) for 1 h. Complex was eluted with buffer containing 500 mM imidazole, and diluted to ∼150 mM NaCl for further purification over an ANX HiTrap ion exchange column (GE Healthcare). The ANX column was washed with ten column volumes of 50 mM Tris-HCl (pH 7.5), 150 mM NaCl, 3 mM TCEP, followed by ten column volumes of 50 mM Bis-Tris (pH 6.0), 150 mM NaCl, 3 mM TCEP, and subjected to a NaCl gradient 150mM-1000m using the Bis-Tris buffer. The CRBN∼DDB1 peak eluted at ∼200 mM NaCl. This peak was collected and further purified by size-exclusion buffer containing 10 mM HEPES pH 7.0, 240 mM NaCl, and 3 mM TCEP. Samples were concentrated to 20mg/mL, aliquoted, flash-frozen, and stored at -80° C.

### Zinc finger protein purification

Domain boundaries for the individually purified MBP (maltose binding protein)-fused zinc finger domains always include the five amino acids N-terminal and one amino acid C-terminal to the zinc finger domains. MBP-fused WT and mutant Ikaros, zinc finger domain proteins were expressed in *E. coli* BL21 (DE3) Star cells (Life Technologies) using 2XYT media (Teknova). Cells were induced at OD_600_ 0.6 for 18 h at 16 °C. Cells were pelleted, frozen, thawed for purification, and resuspended in B-PER Bacterial Protein Extraction buffer (Thermo Fisher) containing 150 µM zinc acetate, 40,000 U benzonase (Novagen), and 1X protease inhibitor cocktail (San Diego Bioscience). Lysates were incubated with amylose resin (NEB) at 4 °C for 1 h before beads were washed. Protein was eluted with buffer containing 200 mM NaCl, 50 mM Tris pH 7.5, 3 mM TCEP, 10% glycerol, 150 µM zinc acetate, and 10 mM maltose. Eluate was concentrated and further purified by size exclusion chromatography over a Superdex 200 16/600 column (GE Healthcare) in buffer containing 200 mM NaCl, 50 mM Tris (pH 7.5), 3 mM TCEP, 10% glycerol, and 150 µM zinc acetate. Cleaved Zinc-finger proteins used in cryo-EM were further as previously described^5^.

### Preparation of samples for cryo-EM

Frozen 20 mg/mL CRBN∼DDB1 complexes were thawed and diluted to 5mg/mL with buffer containing 20 mM HEPES pH 7, 150 mM NaCl, 3mM TCEP. Samples were kept at 4° C throughout the sample preparation process. Immediately prior to plunge-freezing in liquid ethane, samples were diluted 10-fold in the same buffer supplemented with 0.011% Lauryl Maltose-Neopentyl Glycol (LMNG) to improve particle dispersity. Quantifoil 300 mesh R1.2/1.3 UltrAuFoil Holey Gold Films were plasma cleaned for 7 seconds using a Solarus plasma cleaner (Gatan, Inc.) with a 75% nitrogen, 25% oxygen atmosphere at 15 W or glow discharged for 25 seconds with Pelco Easiglow 91000 (Ted Pella, Inc.) in ambient vacuum. To further limit complex dissociation during cryo-EM grid preparation, grids were pre-treated by applying a 4 µL solution containing 10 µM CRBN agnostic mutant Ikaros residues 140-196 Q146A G151N to the surface of the grid ∼1 minute prior to sample application, then blotted from behind with Whatman 1 filter paper. 4 µL of the CRBN∼DDB1 sample was then immediately applied and blot-plunged using a manual plunge freezer in a 4° C cold room with >95% humidity. Grids were blotted for ∼4 s and immediately plunged into a liquid ethane pool cooled by liquid nitrogen. Samples containing CELMoD compounds were prepared the same way, with 10x molar equivalent of drug added 2 minutes prior to dilution and application to the grid, yielding a buffer concentration 20 mM HEPES pH 7, 150 mM NaCl, 3mM TCEP, <1% DMSO. Similarly, Ikaros samples were prepared by incubation of 2:1 Ikaros:CRBN molar excess 2 minutes prior to dilution and application to the grid, or 20:1 excess for the sample with mezigdomide (**Fig. 3C**).

### Cryo-EM data acquisition

Cryo-EM data were collected on a Thermo-Fisher Talos Arctica transmission electron microscope operating at 200 keV using parallel illumination conditions^17^. Micrographs were acquired using a Gatan K2 Summit direct electron detector, operated in electron counting mode applying a total electron exposure of 62.5 e^-^/Å ^2^. The Leginon data collection software^18^ was used to collect micrographs at 36,000x nominal magnification (1.15 Å /pixel at the specimen level) or 45,000x nominal magnification (0.91 Å /pixel at the specimen level) with a nominal defocus set to 0.8 µm-1.2 µm under focus. Stage movement was used to target the center of four 1.2 µm holes for focusing, and an image shift was used to acquire high magnification images in the center of each of either four or sixteen targeted holes. Detailed dataset-specifics are available in **Supplementary Table 1**. In brief, 1,033 micrographs were collected at 1.15 Å pixel for apo CRBN∼DDB1, 2,257 micrographs were collected at 0.91 Å pixel for CRBN∼DDB1 + Pomalidomide, 358 micrographs were collected at 1.15 Å pixel for CRBN∼DDB1 + Iberdomide, 2,474 micrographs were collected at 1.15Å pixel for CRBN∼DDB1+ Iberdomide + Ikaros ZF1-2-3, 3,426 micrographs were collected at 1.15 Å pixel for CRBN∼DDB1 + mezigdomide, and 1,618 micrographs were collected at 1.15 Å pixel for CRBN∼DDB1 + mezigdomide + Ikaros ZF1-2-3.

### Cryo-EM image analysis

A typical processing pipeline is shown in **Supplementary Fig. 5**. Cryo-EM movies were transferred to Warp v1.0.6-v1.0.9^19^ for motion correction, CTF correction, and particle picking. For each dataset, ∼500,000-2,000,000 particles were extracted with 224 pixel box size (∼260Å ^2^), imported to cryoSPARC v2.1-v3.3.2^20^ for liberal cleanup based on 2D classification and all subsequent processing steps unless otherwise noted. Initial models were generated Ab initio from selected particles and used to seed homogeneous or NU-refinement of all quality particles. Classification of BPB position, and CRBN conformation were automatically achieved through iterative heterogeneous classification, 3DVA clustering with standard settings, and 3D classification without alignment using PCA initialization, O-EM learning rate init 0.5, O-EM half life iterations 100, and class similarity 0.2-0.25. Validation or corroborating classification of CRBN∼DDB1 dynamic states was performed in RELION v2.1-v3.1^21^ with single reference 3D classification +/- alignment +/- exclusionary masks focusing on CRBN dynamics or DDB1 dynamics. In total, over 30 datasets were analyzed to improve confidence in global population percentage estimates. Small subsets of particles were chosen as final models, selected for homogeneity across the entire particle, quality of map features, and estimated resolution by FSC at a cutoff of 0.143. Per-particle defocus and up to fourth-order optical aberrations were corrected in cryoSPARC’s NU-refinement protocol. Particles contributing to all three structures containing mezigdomide in **Fig. 3B** were pooled and locally refined with 10° rotation search and 5 Å shift search using a soft mask shown in **Supplementary Fig. 6** to generate the inset.

### Atomic model building and refinement

Model building and refinement were initiated with published models for CRBN∼DDB1, Ikaros, CC-220, and Pomalidomide. Restraints and original PDB for mezigdomide were generated with GRADE server^22^. Iterative rounds of model building and refinement were performed in PHENIX v1.19.2^23^ and Coot 0.9-pre EL revision 8983^24^ until reasonable agreement between the model and data were achieved. In some cases, models were improved with composite maps of focused, local refinements, but validation is performed on non-composite maps (**Supplementary Table 2**). UCSF Chimera and ChimeraX^25^ were used to interpret the EM reconstructions and atomic models, as well as to generate figures.

### *in vitro* ubiquitination assays

Purified E1, E2, ubiquitin, CUL4A–RBX1, CRBN–DDB1, and Ikaros ZF1-2-3 (MBP fusion) proteins were used to reconstitute the ubiquitination of Ikaros ZF1-2-3 in vitro. Human CRBN–DDB1 complex, human CUL4A and RBX1 complex were expressed and purified as previously described^11^. Purified recombinant human Ube1 E1 (E-305), UbcH5a E2 (E2-616), and ubiquitin (U-100H) were purchased from R&D systems. Components were mixed to final concentrations of 1 µM Ube1, 25 µM UbcH5a, 200 µM ubiquitin, 1 µM CUL4–RBX1, 2 µM CRBN–DDB1, and 5 µM MBP-Ikaros-ZF1-2-3, with or without 3 µM pomalidomide, CC-220, and mezigdomide in assay buffer (20 mM HEPES pH 7.5, 150 mM NaCl, 10 mM MgCl_2_). After pre-incubation for 10 min at room temperature, ubiquitination reactions were started by addition of ATP to a final concentration of 10 mM. Reactions were incubated at 30 °C for 30 minutes or 1 hour before separation by SDS–PAGE followed by immunoblot analysis for MBP-substrate proteins (Anti-MBP-probe Antibody (R3.2) (Santa Cruz sc-53366)).

### Hydrogen-deuterium exchange (HDX) detected by mass spectrometry (MS)

#### Peptide Identification

Differential HDX-MS experiments were conducted as previously described with a few modifications^26^. Peptides were identified using tandem MS (MS/MS) with an Orbitrap mass spectrometer (Q Exactive, ThermoFisher). Product ion spectra were acquired in data-dependent mode with the top five most abundant ions selected for the product ion analysis per scan event. The MS/MS data files were submitted to Mascot (Matrix Science) for peptide identification. Peptides included in the HDX analysis peptide set had a MASCOT score greater than 20 and the MS/MS spectra were verified by manual inspection. The MASCOT search was repeated against a decoy (reverse) sequence and ambiguous identifications were ruled out and not included in the HDX peptide set.

#### HDX-MS analysis

5 µl of 10µM CRBN∼DDB1 sample +/- Iberdomide +/- Ikaros ZF1-2-3 was diluted into 20 µl D_2_O buffer (10 mM HEPES, pH 7.0; 240 mM NaCl; 3 mM TCEP) and incubated for various time points (0, 10, 60, 300, and 900 s) at 4°C. The deuterium exchange was then slowed by mixing with 25 µl of cold (4°C) 5 M urea, 50mM TCEP, and 1% trifluoroacetic acid. Quenched samples were immediately injected into the HDX platform. Upon injection, samples were passed through an immobilized pepsin column (2 mm x 2 cm) at 200 µl min^-1^ and the digested peptides were captured on a 2 mm x 1 cm C_8_ trap column (Agilent) and desalted. Peptides were separated across a 2.1 mm x 5 cm C_18_ column (1.9 µl Hypersil Gold, ThermoFisher) with a linear gradient of 4% - 40% CH_3_CN and 0.3% formic acid, over 5 min. Sample handling, protein digestion and peptide separation were conducted at 4°C. Mass spectrometric data were acquired using an Orbitrap mass spectrometer (Exactive, ThermoFisher). HDX analyses were performed in triplicate, with single preparations of each complex form. The intensity weighted mean m/z centroid value of each peptide envelope was calculated and subsequently converted into a percentage of deuterium incorporation. This is accomplished determining the observed averages of the undeuterated and fully deuterated spectra and using the conventional formula described elsewhere^27^. Statistical significance for the differential HDX data is determined by an unpaired t-test for each time point, a procedure that is integrated into the HDX Workbench software^28^. Corrections for back-exchange were made on the basis of an estimated 70% deuterium recovery, and accounting for the known 80% deuterium content of the deuterium exchange buffer.

#### Data Rendering

The HDX data from all overlapping peptides were consolidated to individual amino acid values using a residue averaging approach. Briefly, for each residue, the deuterium incorporation values and peptide lengths from all overlapping peptides were assembled. A weighting function was applied in which shorter peptides were weighted more heavily and longer peptides were weighted less. Each of the weighted deuterium incorporation values were then averaged to produce a single value for each amino acid. The initial two residues of each peptide, as well as prolines, were omitted from the calculations. This approach is similar to that previously described^29^.

## Data Availability

Atomic models have been deposited to the Protein Databank (PDB, https://www.rcsb.org) under accession codes 8cvp, 8d7u, 8d7v, 8d7w, 8d7x, 8d7y, 8d7z, 8d80, 8d81 as described in **Supplementary Table 2**. Electron microscopy reconstructions have been deposited to the Electron Microscopy Databank (EMDB, https://www.emdataresource.org) under accession codes EMDB-27012, EMDB-27234, EMDB-27235, EMDB-27236, EMDB-27237, EMDB-27238, EMDB-27239, EMDB-27240, EMDB-27241, EMDB-27242 as described in **Supplementary Table 1**.

## Acknowledgements

We thank J.C. Ducom at Scripps Research High Performance Computing and C. Bowman at Scripps Research for computational support, as well as B. Anderson at the Scripps Research Electron Microscopy Facility for microscopy support. Computational analyses of EM data were performed using shared instrumentation funded by NIH S10OD021634 to G.C.L.

## Supplementary Materials

**Supplementary Figure 1.**
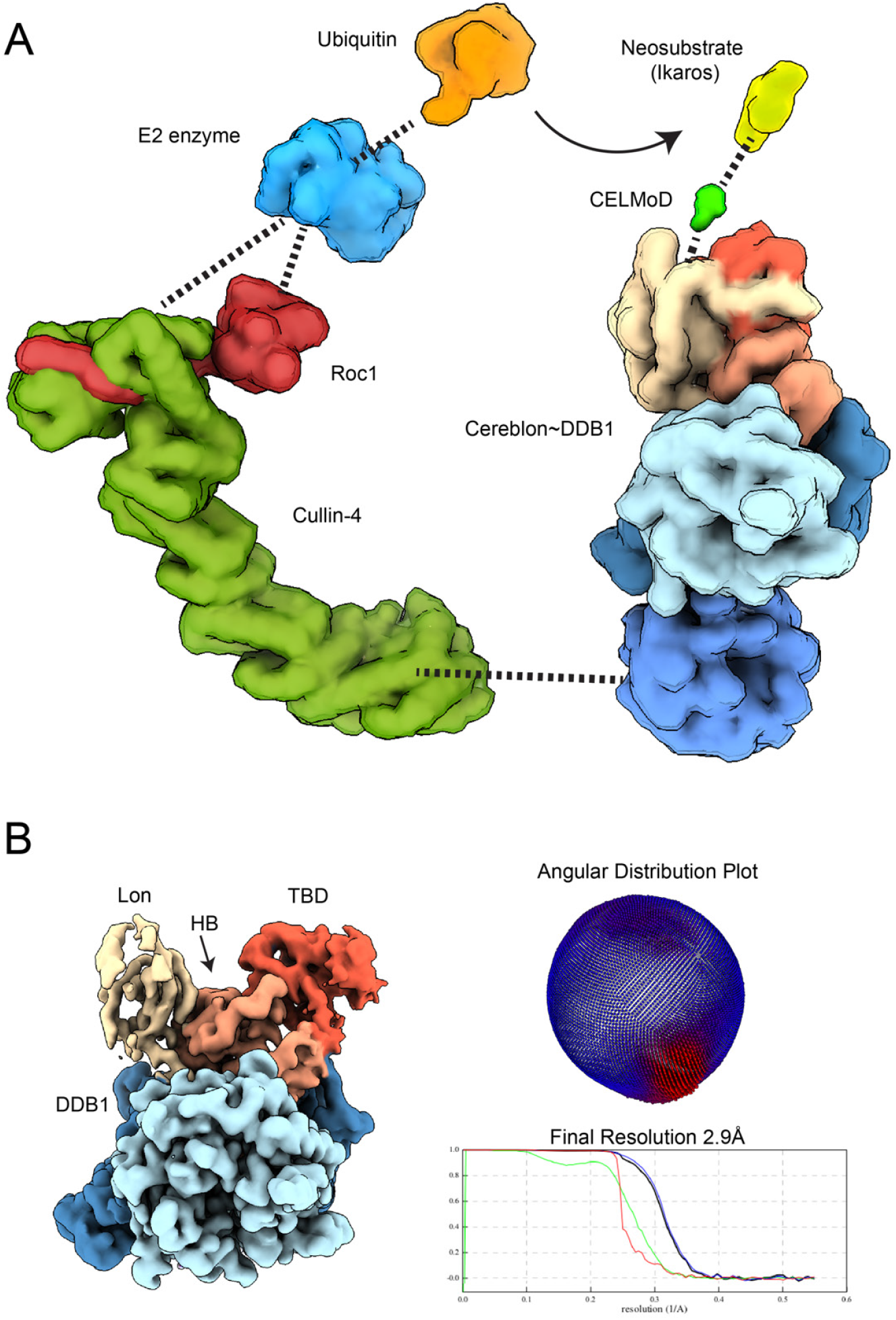
CRBN adopts an open conformation with DDB1^ΔBPB^. **(A)** Schematic diagram of the CRL4∼CRBN complex. At one end, Cul4∼Roc1 recruits a Ubiquitin-loaded E2 and at the other end, coordinates CRBN∼DDB1 which recruits neosubstrates in the presence of CELMoD agents. This scaffold positions Ub for direct transfer to the substrate. **(B)** ∼3 Å resolution unsharpened cryo-EM reconstruction of CRBN∼DDB1^ΔBPB^ in the unliganded apo form. CRBN adopts exclusively the open conformation in absence of CELMoD agents.

**Supplementary Figure 2.**
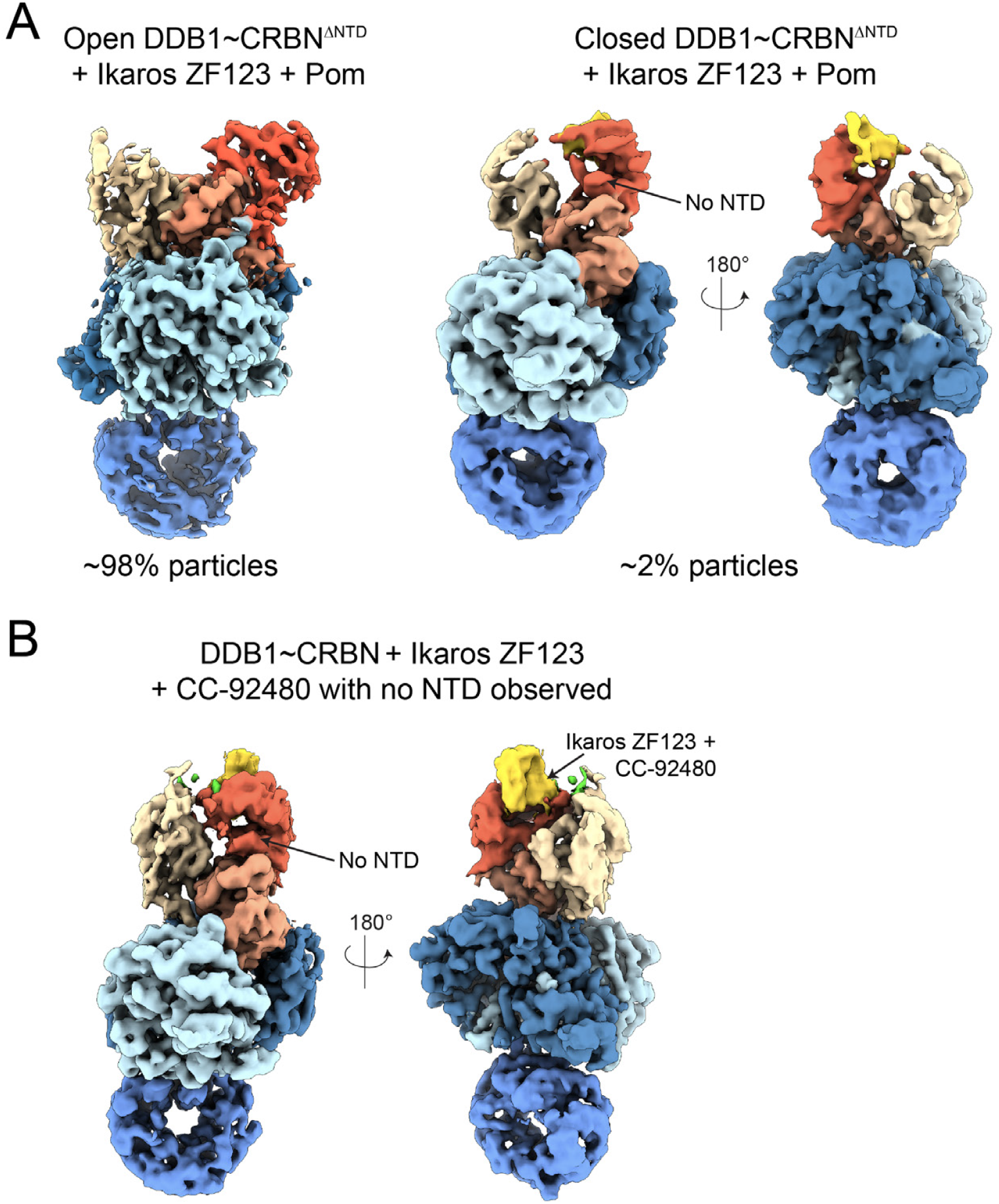
CRBN N-terminal domain. **(A)** >98% of CRBN^ΔNTD^ particles are found in the open conformation in the presence of pomalidomide and Ikaros ZF1-2-3. The 2% of particles adopting CRBN^closed^ have a poorly ordered sensor loop, but are able to bind Ikaros. As expected, no density for the N-terminal seatbelt is observed for this construct. **(B)** In the presence of mezigdomide, CRBN^closed^ with Ikaros ZF1-2-3 bound can be observed without NTD docking, suggesting the CELMoD compound staples CRBN^closed^ and bypasses the requirement for the NTD.

**Supplementary Figure 3.**
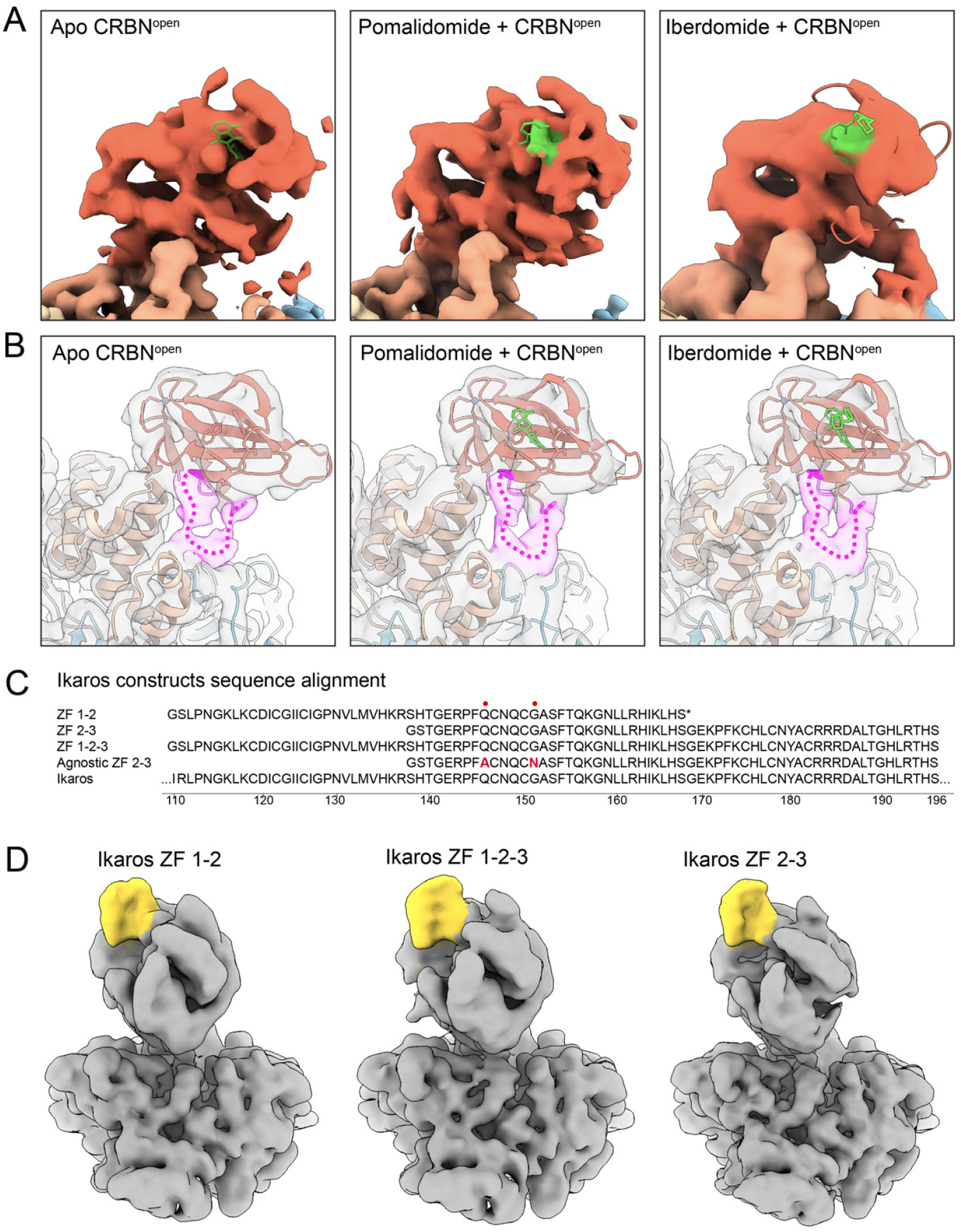
Pomalidomide is recruited to both CRBN^open^ and CRBN^closed^ but recruits only a single ZF to CRBN^closed^. **(A)** View of TBD (red) from low-resolution cryo-EM reconstructions of unliganded apo CRBN^open^, pomalidomide-bound CRBN^open^, and iberdomide-bound CRBN^open^ (compounds colored green) showing that the tri-Trp binding pocket drug is conspicuously empty in the absence of compound (semi-transparent pomalidomide indicates the location of the binding pocket), but is able to efficiently recruit compounds to the open conformation with high occupancy. TBD∼pomalidomide and TBD∼CC-220 from PDB 6h0g and PDB 5v3o, respectively, are shown based on rigid-body docking of the PDBs into the cryo-EM densities. **(B)** Unsharpened maps from (A), shown as in Fig. 1A, with common indication for the sensor loop backbone in purple. **(C)** Sequence alignment for Ikaros proteins that were used in these studies. Ikaros comprises multiple zinc finger motifs in tandem, here we have utilized ZF1-2, ZF2-3, ZF1-2-3, and a CRBN-agnostic mutated ZF2-3 for preparation of cryo-EM grids. **(D)** Unsharpened volumes of ∼4 Å cryo-EM reconstructions of CRBN∼DDB1^ΔBPB^ bound to different versions of Ikaros tandem ZF protein. Only a single zinc finger motif (colored gold) is observed to bind to the TBD. Other zinc finger domains are not visible, presumably because they are flexibly attached and do not bind CRBN.

**Supplementary Figure 4.**
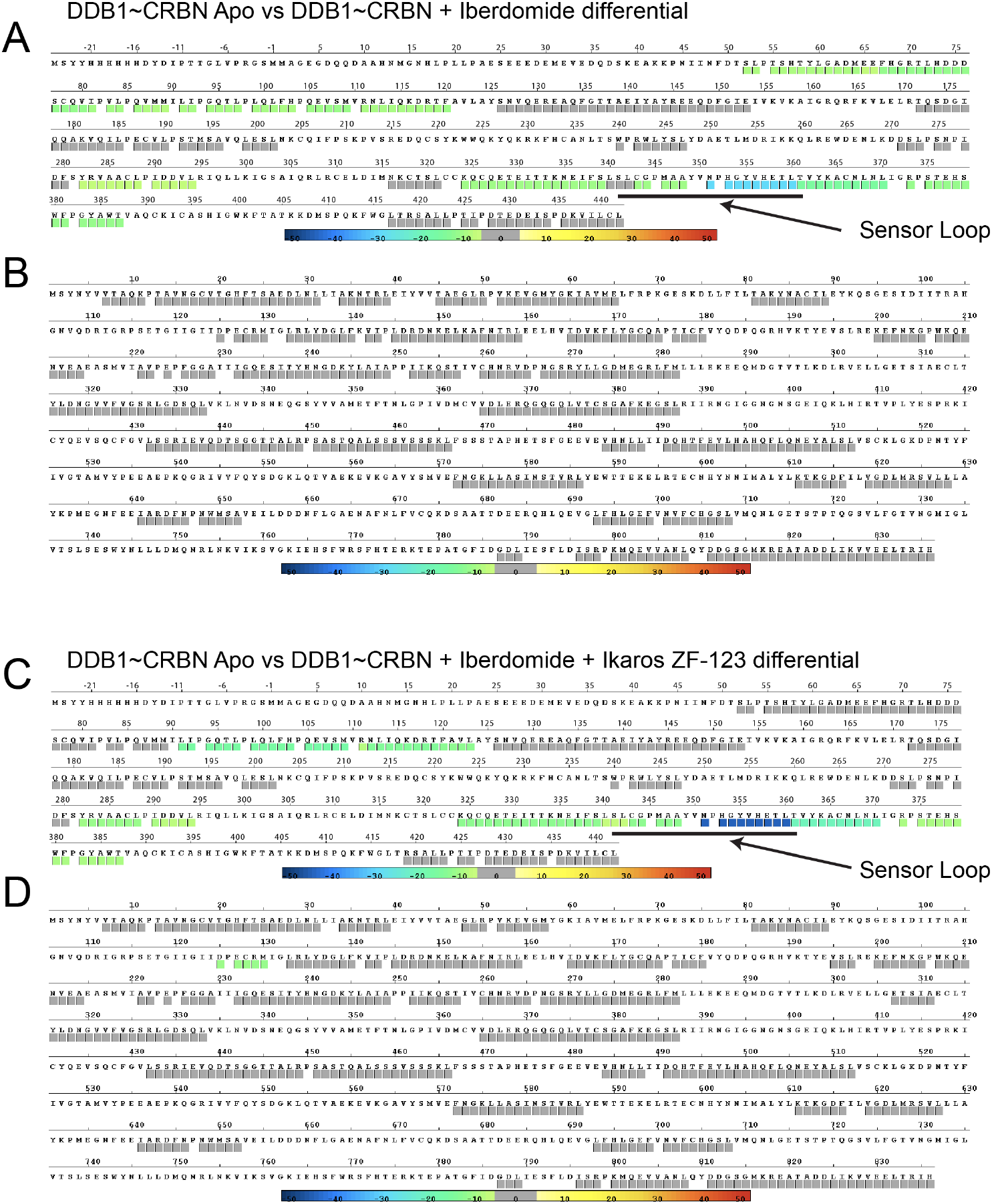
Hydrogen-deuterium exchange consolidated view. **(A)** Per-residue peptide mapping of CRBN HDX differential upon addition of iberdomide to CRBN∼DDB1. Colored boxes underneath the amino acid sequence denote digested peptides detected in mass spectrometry experiments. Regions with no change in exchange upon the addition of iberdomide have grey boxes while regions with reduced D_2_O exchange are colored according to the scale. **(B)** Per-residue peptide mapping of DDB1 HDX differential upon addition of iberdomide to CRBN∼DDB1. **(C)** Per-residue peptide mapping of CRBN HDX differential upon addition of iberdomide and Ikaros ZF1-2-3 to CRBN∼DDB1. **(D)** Per-residue peptide mapping of DDB1 HDX differential upon addition of iberdomide and Ikaros ZF1-2-3 to CRBN∼DDB1.

**Supplementary Figure 5.**
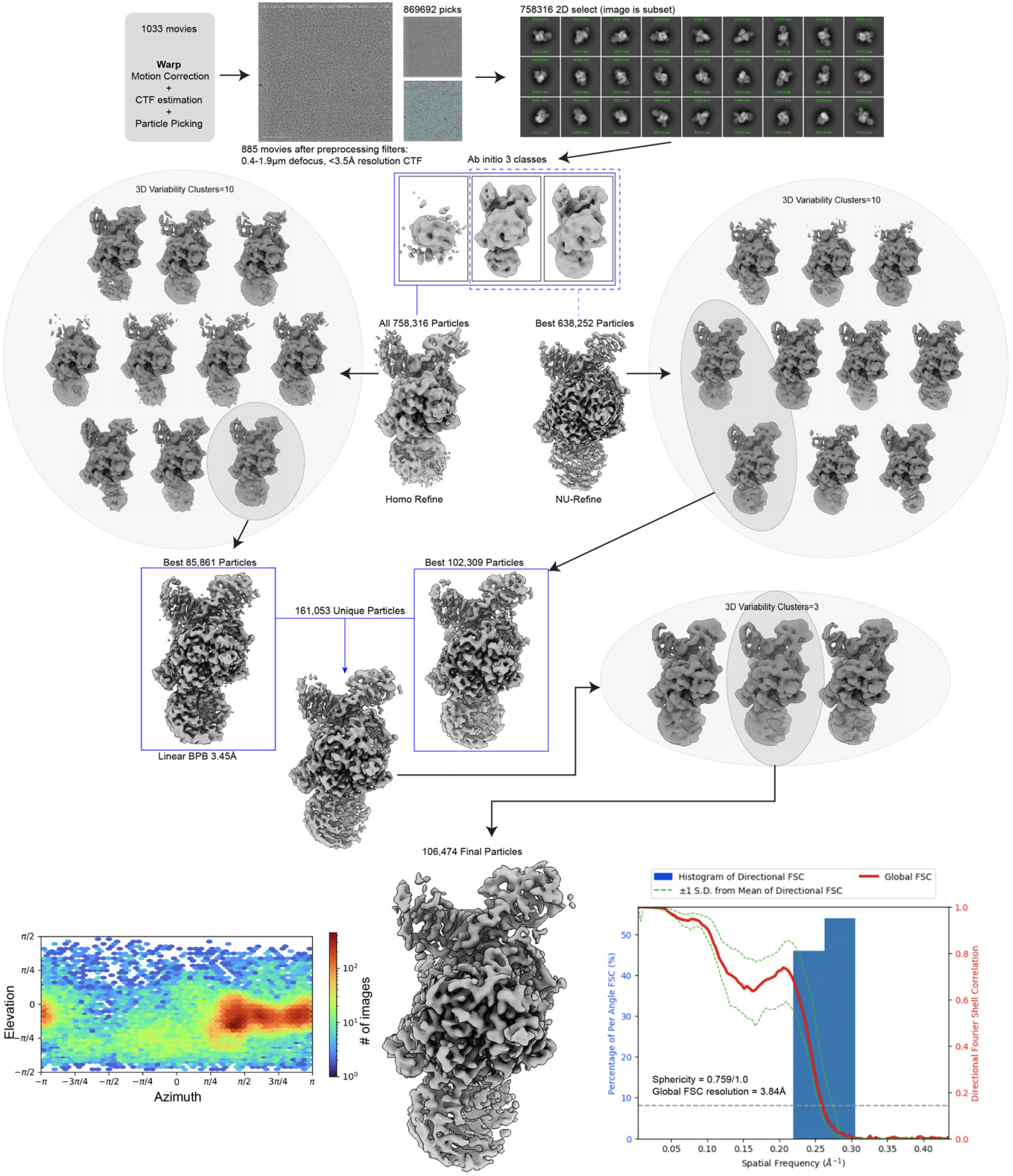
Processing workflow. Representative processing workflow described for CRBN∼DDB1 in the unliganded apo form. Final reconstruction is shown at the bottom centered between angular distribution plot and 3DFSC^30^.

**Supplementary Figure 6.**
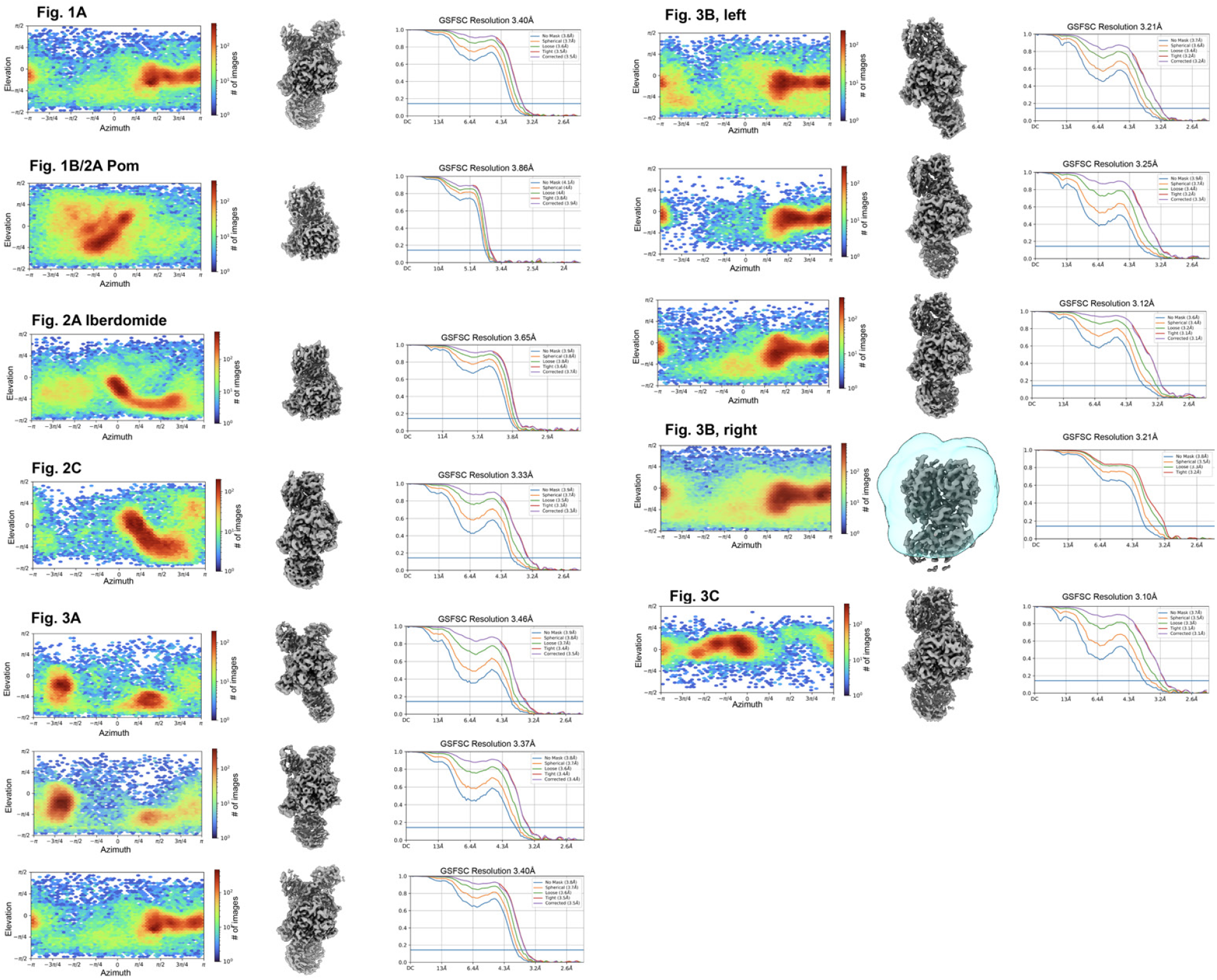
Gallery of Euler distribution and FSCs of structures. Gallery of structures reported in this manuscript are shown between their angular distribution plots and reported resolutions via gold-standard FSC.

**Supplementary Table 1.**
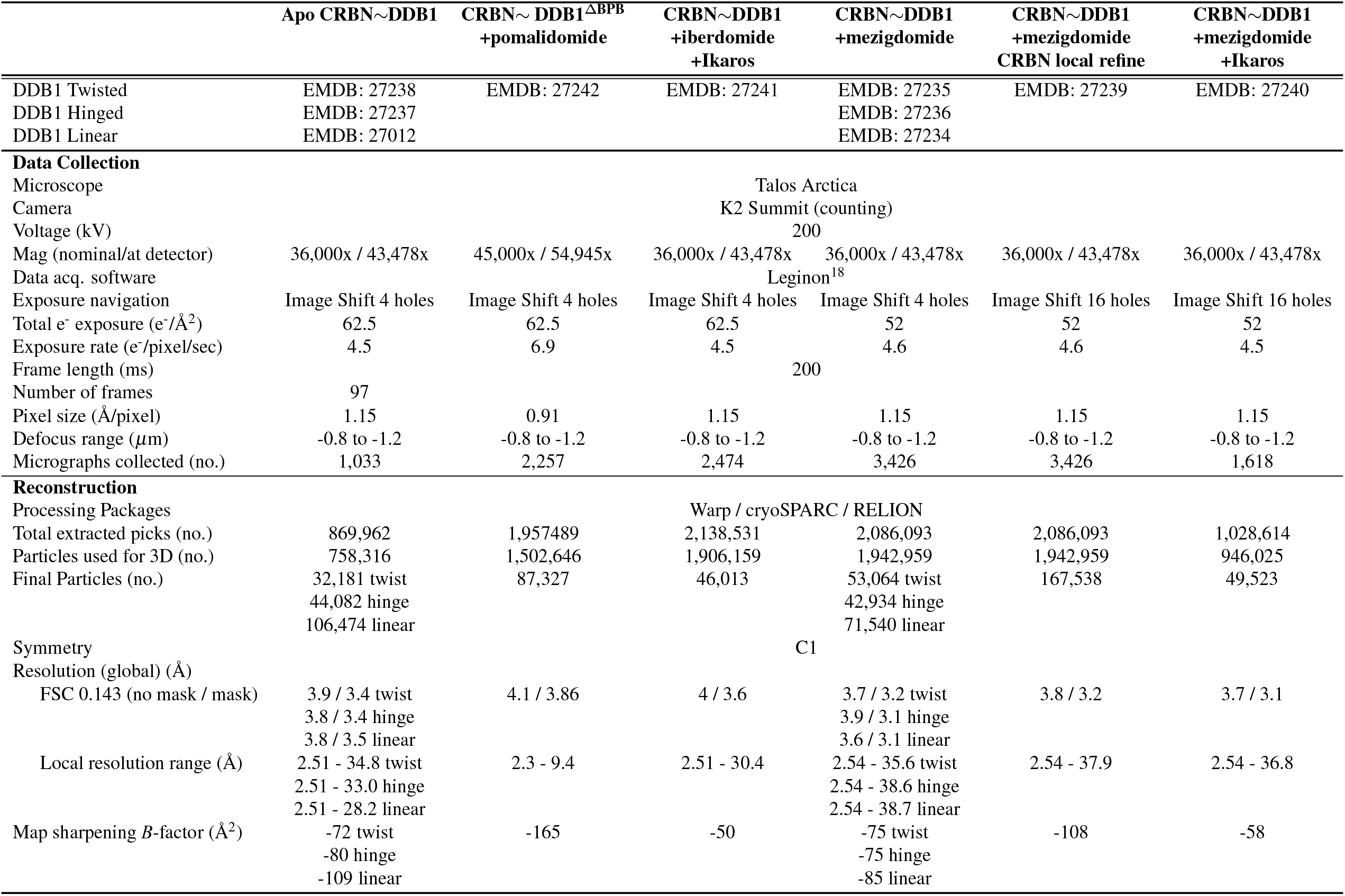
Cryo-EM data collection and image analysis.

|  | Apo CRBN~DDB1 | CRBN~DDB1<sup>ΔBPB</sup><br>+pomalidomide | CRBN~DDB1<br>+iberdomide<br>+Ikaros | CRBN~DDB1<br>+mezigdomide | CRBN~DDB1<br>+mezigdomide<br>CRBN local refine | CRBN~DDB1<br>+mezigdomide<br>+Ikaros |
| --- | --- | --- | --- | --- | --- | --- |
| DDB1 Twisted | EMDB: 27238 | EMDB: 27242 | EMDB: 27241 | EMDB: 27235 | EMDB: 27239 | EMDB: 27240 |
| DDB1 Hinged | EMDB: 27237 |  |  | EMDB: 27236 |  |  |
| DDB1 Linear | EMDB: 27012 |  |  | EMDB: 27234 |  |  |
| **Data Collection** |  |  |  |  |  |  |
| Microscope |  |  |  | Talos Arctica |  |  |
| Camera |  |  |  | K2 Summit (counting) |  |  |
| Voltage (kV) |  |  |  | 200 |  |  |
| Mag (nominal/at detector) | 36,000x / 43,478x | 45,000x / 54,945x | 36,000x / 43,478x | 36,000x / 43,478x | 36,000x / 43,478x | 36,000x / 43,478x |
| Data acq. software |  |  |  | Leginon<sup>18</sup> |  |  |
| Exposure navigation | Image Shift 4 holes | Image Shift 4 holes | Image Shift 4 holes | Image Shift 4 holes | Image Shift 16 holes | Image Shift 16 holes |
| Total e<sup>-</sup> exposure (e<sup>-</sup>/Å<sup>2</sup>) | 62.5 | 62.5 | 62.5 | 52 | 52 | 52 |
| Exposure rate (e<sup>-</sup>/pixel/sec) | 4.5 | 6.9 | 4.5 | 4.6 | 4.6 | 4.5 |
| Frame length (ms) |  |  |  | 200 |  |  |
| Number of frames | 97 |  |  |  |  |  |
| Pixel size (Å/pixel) | 1.15 | 0.91 | 1.15 | 1.15 | 1.15 | 1.15 |
| Defocus range (μm) | -0.8 to -1.2 | -0.8 to -1.2 | -0.8 to -1.2 | -0.8 to -1.2 | -0.8 to -1.2 | -0.8 to -1.2 |
| Micrographs collected (no.) | 1,033 | 2,257 | 2,474 | 3,426 | 3,426 | 1,618 |
| **Reconstruction** |  |  |  |  |  |  |
| Processing Packages |  |  |  | Warp / cryoSPARC / RELION |  |  |
| Total extracted picks (no.) | 869,962 | 1,957,489 | 2,138,531 | 2,086,093 | 2,086,093 | 1,028,614 |
| Particles used for 3D (no.) | 758,316 | 1,502,646 | 1,906,159 | 1,942,959 | 1,942,959 | 946,025 |
| Final Particles (no.) | 32,181 twist<br>44,082 hinge<br>106,474 linear | 87,327 | 46,013 | 53,064 twist<br>42,934 hinge<br>71,540 linear | 167,538 | 49,523 |
| Symmetry |  |  |  | C1 |  |  |
| Resolution (global) (Å) |  |  |  |  |  |  |
| FSC 0.143 (no mask / mask) | 3.9 / 3.4 twist<br>3.8 / 3.4 hinge<br>3.8 / 3.5 linear | 4.1 / 3.86 | 4 / 3.6 | 3.7 / 3.2 twist<br>3.9 / 3.1 hinge<br>3.6 / 3.1 linear | 3.8 / 3.2 | 3.7 / 3.1 |
| Local resolution range (Å) | 2.51 - 34.8 twist<br>2.51 - 33.0 hinge<br>2.51 - 28.2 linear | 2.3 - 9.4 | 2.51 - 30.4 | 2.54 - 35.6 twist<br>2.54 - 38.6 hinge<br>2.54 - 38.7 linear | 2.54 - 37.9 | 2.54 - 36.8 |
| Map sharpening *B*-factor (Å<sup>2</sup>) | -72 twist<br>-80 hinge<br>-109 linear | -165 | -50 | -75 twist<br>-75 hinge<br>-85 linear | -108 | -58 |

**Supplementary Table 2.**
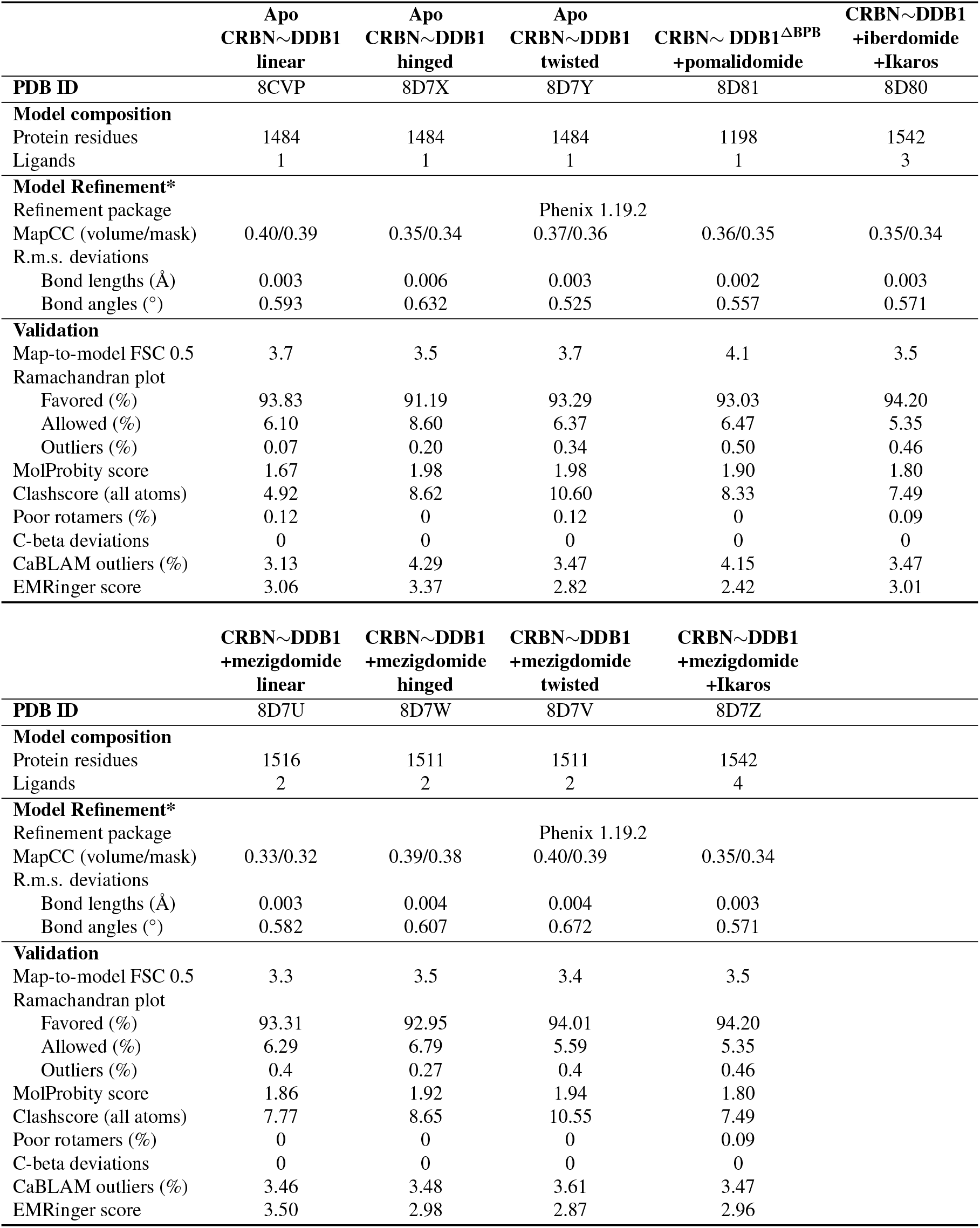
Atomic modeling, refinement, and validation statistics.

